# Modulating NLRP3 splicing with antisense oligonucleotides to control pathological inflammation

**DOI:** 10.1101/2024.09.06.611206

**Authors:** Roni Klein, Janset Onyuru, Jessica L. Centa, Estela M. Viera, Fox J. Duelli, Christopher D. Putnam, Hal M. Hoffman, Michelle L. Hastings

**Affiliations:** Center for Genetic Diseases, Chicago Medical School, Rosalind Franklin University of Medicine and Science, North Chicago, Illinois, USA; School of Graduate and Postdoctoral Studies, Rosalind Franklin University of Medicine and Science, North Chicago, Illinois, USA; Department of Pediatrics, University of California San Diego, La Jolla, California, USA; Department of Pharmacology, University of Michigan Medical School, Ann Arbor, Michigan, USA; Department of Medicine, University of California San Diego, La Jolla, California, USA

## Abstract

Inflammation has an essential role in healing. However, over-active inflammation disrupts normal cellular functions and can be life-threatening when not resolved. The NLR family pyrin domain-containing 3 (NLRP3) inflammasome, a component of the innate immune system, is an intracellular multiprotein complex that senses stress-associated signals, and, for this reason is a promising therapeutic target for treating unresolved, pathogenic inflammation. Alternative splicing of *NLRP3* RNA has been suggested as a regulatory mechanism for inflammasome activation, as some spliced isoforms encode NLRP3 proteins with compromised function. Here, we take advantage of this natural regulatory mechanism and devise a way to control pathogenic inflammation using splice-switching antisense oligonucleotides (ASOs). To identify and induce NLRP3 isoforms lacking inflammatory activity, we tested a series of ASOs, each targeting a different exon, to determine the most effective strategy for down-regulating NLRP3. We identify several ASOs that modulate *NLRP3* splicing, reduce NLRP3 protein, and decrease inflammasome signaling in vitro. The most effective ASO suppresses systemic inflammation in vivo in mouse models of acute inflammation and cryopyrin-associated periodic syndrome (CAPS). Overall, these results demonstrate how ASOs can be used to systematically engineer proteins with modified functions and treat pathological inflammation in mice by reducing functional NLRP3.

## Introduction

The innate immune system defends against infectious and non-infectious threats, and inflammasomes are crucial mediators of this response. These intracellular multiprotein complexes detect microbial motifs and danger-associated molecules and mount an inflammatory response (1). Inflammasomes are categorized by their structural motifs. NLR-type inflammasomes have receptors with nucleotide-binding leucine-rich repeat (NLR) domains (2). The NLRP3 inflammasome (NLR family pyrin-containing protein 3) regulates the cellular response to pathogens and inflammatory insults that threaten cells. Once activated, NLRP3 forms an inflammasome complex that includes ASC (apoptosis-associated speck-like protein containing a CARD) and caspase 1 (CASP1), which self-cleaves and mediates interleukin-1β (IL-1β, IL1B) and interleukin-18 (IL-18, IL18) maturation and secretion. Mature CASP1 also disinhibits gasdermin D (GSDMD), which leads to pore formation on the cell membrane and pyroptotic cell death, thereby eliminating infected or damaged cells (3). If not resolved, the pro-inflammatory cytokine cascade can alter normal cellular functions and further propagate an inflammatory response that can lead to tissue damage (4).

Although inflammation is a natural way for the body to respond to pathogens and eliminate damaged cells, endogenous stress signals can lead to aberrant NLRP3-mediated inflammation (5–8). Persistent NLRP3 inflammasome activation has been implicated in the pathogenesis of autoinflammatory, neurodegenerative, and metabolic diseases (9–11). Furthermore, gain of function mutations in *NLRP3* cause cryopyrin-associated periodic syndrome (CAPS), a rare pediatric autoinflammatory disease of varying severity resulting from dysregulation of the NLRP3 inflammasome and uncontrolled release of inflammatory cytokines such as IL-β and IL-18 and pyroptosis (12, 13). In its most severe forms, Muckle-Wells syndrome (MWS) and neonatal-onset multisystem inflammatory disorder (NOMID), the disease presents with systemic, cutaneous, musculoskeletal, and central nervous system (CNS) inflammation, leading to progressive organ damage and dysfunction (14). Inhibition of inflammasome signaling has been a therapeutic goal in cases of chronic and pathogenic inflammation that becomes detrimental to life.

A number of approaches are being pursued in the clinic to therapeutically block the inflammasome; though, to date, there are still no good treatment options for many forms of NLRP3-mediated inflammation (3, 11). An alternative therapeutic approach is modulation of NLRP3 expression with antisense oligonucleotides (ASOs). ASOs are short, synthetic, modified nucleic acids that can be designed to specifically base-pair to a target RNA and modulate its expression. Splice-switching ASOs are designed to alter splicing by base-pairing to and blocking recognition of RNA sequences that are important for proper splicing (15). In this way, ASOs can be used as a therapeutic approach for modulating splicing to increase or decrease mRNA transcripts encoding protein isoforms with differential activities. ASOs have many favorable drug-like qualities, including high target specificity, stability in cells and favorable safety profiles (16, 17). Indeed, ASO medicines are having clinical success for a variety of conditions (15, 18).

To investigate the potential of splice-switching ASOs for NLRP3-associated pathogenic inflammation, we systematically evaluated the activity of NLRP3 isoforms induced by ASO-mediated splice-switching of each exon and identified ASOs that effectively suppress NLRP3-dependent activation in vitro and in vivo. Treatment with our most effective ASO partially suppresses systemic inflammation in a mouse model of acute inflammation and prolongs survival in a murine model of CAPS, demonstrating that modulation of *NLRP3* with splice-switching ASOs may have therapeutic value in treating NLRP3-driven inflammatory disease.

## Methods

### Antisense oligonucleotides and nomenclature

Splice switching antisense oligonucleotides are phosphorodiamidate morpholinos (PMO) (Gene Tools, LLC) (Supplemental Table 1). A non-targeting PMO was used as a negative control (Gene Tools, standard control oligo). Lyophilized ASOs were formulated in filtered deionized H_2_O, 0.9% saline, or Dulbecco’s phosphate-buffered saline (DPBS). The identity of ASO-induced *NLRP3* isoforms was determined by Sanger sequencing. Analysis of potential off-target base-pairing of human and mouse ASO-Δ2 (gggenome.dbcls.jp), found no other matches in the human or mouse genomes that had fewer than four (human) or three (mouse) nucleotide mismatches and no more than 13 (human) or 11 (mouse) contiguous nucleotides.

### Cell culture and transfections

THP-1 cells (ATCC TIB-202), a monocytic cell line isolated from peripheral blood from an acute monocytic leukemia patient, were cultured in Roswell Park Memorial Institute (RPMI) 1640 supplemented with 10% fetal bovine serum (FBS) and 0.05 mM 2-mercaptoethanol at 37 °C and 5% carbon dioxide (CO_2_). THP-1 cells were seeded at 5×10^5^ cells/mL, differentiated with 50 ng/mL phorbol 12-myristate 13-acetate (PMA; Sigma-Aldrich) for 24 hours, and transfected 18 hours later (44, 45). Immortalized murine bone-marrow derived macrophages (iBMDMs) (a gift from Dr. Venkat Magupalli and Dr. Hao Wu) were cultivated in Dulbecco’s Modified Eagle’s medium supplemented with 10% FBS at 37 °C and 5% CO_2_. iBMDMs were plated at 40-50% confluency and incubated for 18 hours prior to transfection. ASOs with a PMO chemistry were transfected into THP-1 and iBMDM at a final concentration of 40 µM, or for the dose responses from 2.5 µM to 40 µM using Endo-Porter (Gene Tools, LLC) for 48 hours according to manufacturer instructions (Gene Tools, LLC).

Human monocyte-derived macrophages (hMDMs) from patients with CAPS harboring an *NLRP3* L353P mutation were separated by Percoll-gradient (Cytiva) centrifugation and subsequently differentiated with 20 ng/mL macrophage colony-stimulating factor (M-CSF, R&D Systems) for 7 days. On day six, cells were transfected with ASO-Δ2 or ASO-C (40 µM final concentration) for 24 hours using Endo-Porter (Gene Tools LLC). Cells were then stimulated with 200 ng/mL LPS (*Escherichia coli* 0111:B4, InvivoGen) for ∼16 hours and supernatants were collected for IL-1β ELISA analysis. The lysate was collected for splicing analysis.

### Inflammasome activation assay

PMA-differentiated THP-1 cells and iBMDMs were transfected with ASOs as described above and incubated for 48 hours. Cells were then primed with 100 ng/mL LPS (*Escherichia coli* 055:B5, Sigma-Aldrich) in supplemented media for 3 hours. Media was subsequently removed and cells were activated with 5 mM ATP (Invivogen) in OptiMem (Thermo Fisher Scientific) for 1 hour. Cells in OptiMem (Thermo Fisher Scientific) were treated with 100 nM MCC950 (Invivogen) for 30 minutes prior to activation with 5 mM ATP for 1 hour. Media was collected for analysis of released signaling molecules and lysate was collected for RNA and protein analysis.

### RNA isolation and analysis

Total RNA was extracted from cells or mouse tissue using TRIzol according to the manufacturer’s instructions (Thermo Fisher Scientific). RNA (1μg) was reverse transcribed using GoScript reverse transcriptase (Promega) and oligo-dT primer. RT-PCR and radiolabeled RT-PCR of cDNA was performed using GoTaq Green (Promega) with or without α-^32^P-deoxycytidine triphosphate, and primers for *NLRP3* (Supplemental Table 1, Figures 1 and 4). PCR products were resolved on either a 2% agarose gel stained with ethidium bromide or on 6% non-denaturing polyacrylamide gels and quantified using Image J or a Typhoon FLA 7000 phosphorimager (GE Healthcare), respectively. The percent of the *NLRP3* mRNA including the targeted exon was calculated relative to the abundance of the predominant ASO-induced isoform after the signal was corrected for the amplicon size or for the number of cytosine nucleotides for agarose and polyacrylamide gels, respectively. Predominant ASO-induced isoforms were isolated from agarose gels, column purified (Cytiva), and Sanger sequenced with *NLRP3* primers (Supplemental Table 1).

**Figure 1.**
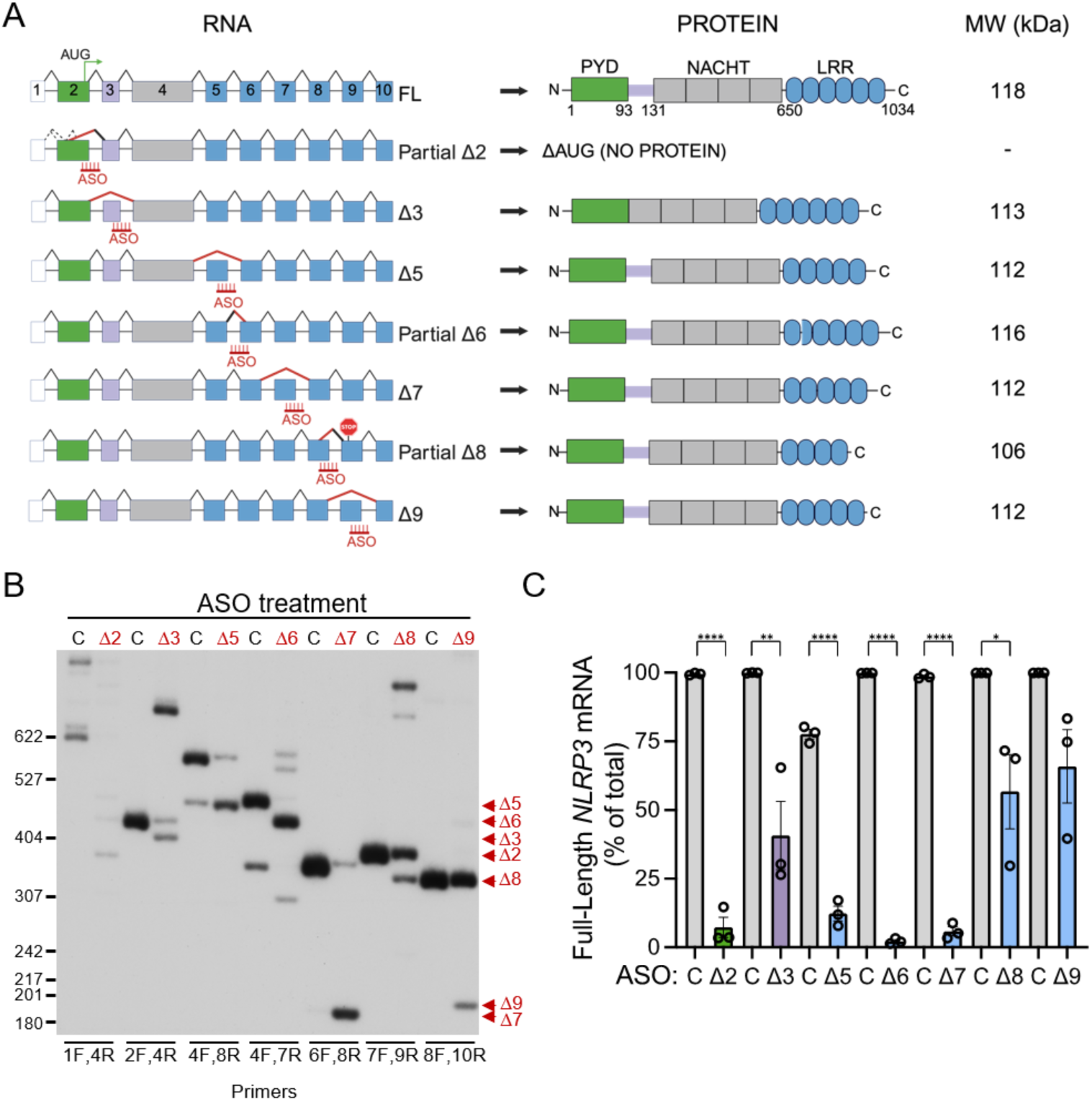
Splice-switching ASOs modulate *NLRP3* alternative splicing. **(A)** Schematic of *NLRP3* spliced and protein isoforms. (left) Full-length (FL) *NLRP3* mRNA and ASO-induced splice variants. Boxes are exons and lines are introns. ASOs are shown in red. Diagonal black lines indicate splicing of introns. Dashed diagonal lines represent natural alternative splicing. Red diagonal lines indicate ASO-induced splicing event. (right) NLRP3 protein domains and predicted protein products translated from the induced spliced isoforms with molecular weights (MW). **(B)** Representative radioactive RT-PCR analysis of *NLRP3* RNA from THP-1 cells treated with the indicated ASOs (40 µM), including a non-targeted ASO control (C), followed by activation with LPS and ATP. Primer sets used to detect the specific ASO-induced splicing event are shown at bottom of gel image. Products were separated by PAGE. Red arrowheads indicate predominant ASO-induced spliced isoform for each ASO. Some ASOs also induced a low level of other alternatively spliced products. **(C)** Quantification of native spliced products (full-length) from *NLRP3* mRNA relative to the predominant ASO-mediated isoforms. Data are presented as mean ± SEM, unpaired t-test compared to matched control, *P<0.05, **P<0.01, ****P<0.0001; n=3 across three independent experiments.

### Immunoblot analysis

Cells were lysed in RIPA buffer (150 mM NaCl, 50 mM Tris-Cl pH 7.6, 1% NP-40, 0.5% sodium deoxycholate, 0.1% SDS) containing protease and phosphatase inhibitor cocktail (Thermo Fisher Scientific). Lysates were centrifuged at 10,000 x g for 20 min at 4 °C and protein concentration was determined by Bradford or Pierce BCA protein assay (Thermo Fisher Scientific). Supernatant was centrifuged using 10 kDa Amicon Ultra Centrifugal Filters (Millipore). Lysates and supernatants were prepared with 4x Bolt lithium dodecyl sulfate (Thermo Fisher Scientifc) and 10x dithiothreitol and heated at 95 °C for 5 minutes. Cell lysates were separated by 4–15% SDS-polyacrylamide electrophoresis (PAGE) Tris-glycine precast gels (Bio-Rad) and supernatants were separated by 12% SDS-PAGE Tris-glycine gels and both were transferred to a 0.45 μm polyvinylidene difluoride blotting membrane (Immobilon-FL, Millipore) at 75V for 1 hour. Membranes were blocked with 5% nonfat dry milk in Tris-buffered saline supplemented with 0.1% Tween 20 and probed with primary antibodies overnight at 4 °C using rabbit NLRP3 (1:1000, D4D8T-15101, Cell Signaling), mouse caspase 1 (1:1000, Casper-1 AG-20B-0042, Adipogen). Membrane was incubated with rabbit α-tubulin (1:1,000, 11224-1-AP, Proteintech) for 1 hour at room temperature. Horseradish peroxidase-conjugated rabbit and mouse secondary antibodies (1:5,000-1:20,000, 32230 and 32260, Thermo Fisher Scientific) were incubated with the membrane for 1 hour at room temperature. Immunoblots were developed using Classico (Immobilon) or SuperSignal West Femto (Thermo Fisher Scientific) substrate and exposed to film. Western blot bands were quantified using ImageJ software.

### ELISA and LDH assay

Supernatant, serum, plasma, and tissue were analyzed for IL-1β (human DY200, mouse DY401), IL-6 (mouse DY406), IL-18 (mouse DY7625), and TNF-ɑ (human DY210, mouse DY410) by ELISA (R&D Systems) according to manufacturer’s instructions. Optical density (450 nm – 540 nm) values are normalized to LPS-and ATP-stimulated cells and utilized to extrapolate the concentration from a standard curve in vivo. LDH release was measured with a CytoTox 96® Non-Radioactive Cytotoxicity Assay (G1780, Promega) and the percent of LDH was calculated relative to LPS and ATP stimulated cells.

### Mice

Mice were maintained at 23 °C with a 12 h light/dark cycle with food and water available *ad libitum*. Wildtype C57BL/6 mice were purchased from The Jackson Laboratory and bred at RFUMS. Mice bearing an aspartate 301 to asparagine (D301N) substitution were generated as previously described (The Jackson Laboratory: B6N.129-*Nlrp3D301NneoR*/J, strain no. 017971) (21, 46). The D301N point mutation results in a conformational change that leads to a ligand-independent constitutive activation of the mutant NLRP3 inflammasome. Due to the presence of an intronic-floxed neomycin resistance cassette, *Nlrp3* gene expression is abolished. Expression of the mutant allele is only achieved when the *Nlrp3* knock-in mice are bred with mice expressing Cre recombinase. *Nlrp3* knock-in mice were bred with mice expressing Cre recombinase under the control of lysozyme promoter (CreL; The Jackson Laboratory, strain no. 004781, B6.129P2-*Lyz2tm1(cre)Ifo*/J), resulting in constitutive activation of mutant *Nlrp3* in the myeloid lineage.

### LPS challenge

Wildtype C57BL/6 female mice (7-8 weeks old) were intraperitoneally (I.P.) injected with ASO-Δ2 (100 mg/kg), ASO-C (100 mg/kg), or PBS on days 1,3, and 6. On day 7, mice were I.P. injected at 7 am with 20 mg/kg ultrapure LPS from E. coli 0111:B4 (Invivogen, tlrl-3pelps) or PBS. Three hours later, the treated mice were euthanized, blood was collected in EDTA-coated tubes through cardiac puncture. For all but one mouse, blood was diluted 1:1 in PBS and added to Ficoll and centrifuged at 400 x g for 30 minutes. For one mouse, blood was collected in EDTA tube and centrifuged 2000 x g for 10 minutes. Plasma was collected and stored in -80 °C until analysis.

### CAPS mice

*Nlrp3*^D301N/+ LysMCre+^ male and female mice were injected with ASO-Δ2 (100 mg/kg) or PBS by subcutaneous injection on P1, P3 and then every 3 days until P24. Mice were monitored daily for growth and survival. A group of mice were euthanized at P12 and tissue and serum were collected. Tissue was snap frozen in liquid nitrogen, and stored in -80 °C until further analysis.

### Histological staining analysis

Mouse skin lesions from the nape were collected following euthanization. Hair around the lesion was removed using hair removal cream (Nair) with a gauze-tipped applicator. After 24 hours of fixation in 10% Buffered Formalin, the paraffin-embedded skin lesions were stained with hematoxylin and eosin (H&E). The slides were then reviewed by a blinded pathologist to identify the dermis and neutrophil infiltration. Scans of all sections were taken using an Olympus VS200 Slide Scanner (UCSD Neuroscience Microscopy Core) at 40X magnification. The number of neutrophils in a single section from each mouse were quantified using a cell classifier on Qupath v0.5.1.

### Structure modeling

Models of the human NLRP3 Δ6 and Δ8 isoforms were initially generated from the predicted protein sequences by Alphafold2 (47). The folds of the individual domains of the resulting models matched that of the experimental NLRP3 structures; however, the resulting conformation did not precisely match that of the known active or inactive conformations (25, 48), which are related by a rigid body motion between the FISNA-NBD-HD1 N-terminal subdomains of the NACHT domain and the WH-HD2-LRR domains (11). To build models of the Δ6 and Δ8 isoforms in the known active and inactive conformations, the Alphafold2-predicted Δ6 and Δ8 leucine-rich repeats were superimposed onto the LRR repeats of the full-length active conformation (PDB id 8ej4) (48) and the full-length inactive conformation (PDB id 7pzc) (25). This analysis was performed by first calculating the affine transformation matrices to superimpose residues 651-683 of the Alphafold2 models onto residues 651-683 for each of the NLRP3 monomers in the inactive and active assemblies; this residue range includes NLRP3 regions involved in the WH-HD2-LRR rigid body motion but avoids residues deleted in the Δ6 and Δ8 isoforms. The affine transformations were then applied to the entire Alphafold2 modeled Δ6 and Δ8 LRR domains (residues 651-1034) and the transformed LRR domains were merged with residues 1-650 of the experimental structures.

### Statistical analysis

All data were analyzed GraphPad Prism (version 10.2.3) and presented as the mean ±SEM. A P value less than 0.05 was considered significant (*P<0.05, **P<0.01, ***P<0.001, ****P<0.0001). All testable data had normal distributions as determined using the D’Agostino & Pearson test for normality. For comparison of two groups, we used the two-tailed unpaired t-test or one-sample t-test. For more than two groups, we utilized one-way or repeated measures ANOVA followed by post-hoc analysis as detailed in figure legends. ASO potency was calculated using the half maximal inhibitory concentration (IC50) after plotting the data on a nonlinear regression curve with a standard slope. Survival curve comparison was analyzed by Gehan-Breslow-Wilcoxon test. The specific statistical test used for each experiment is specified in the figure legends.

### Study approval

All protocols met ethical standards for animal experimentation and were approved by the Institutional Animal Care and Use Committee of Rosalind Franklin University of Medicine and Science and University of California San Diego. Studies performed using PBMCs obtained from patients with CAPS received the approval of the University of California Human Research Protection Program committee, and informed consent was obtained from the subjects before the study.

### Data availability

Values for all data points in graphs are reported in the Supporting Data Values file. Uncropped RT-PCR gel scans and western blots are provided in a supplemental file. Any other data can be requested from the corresponding author.

## Results

### Systematic ASO screening reveals induced NLRP3 spliced isoforms that suppress inflammasome activation

To identify the most effective approach to reducing NLRP3 activity using splice-switching ASOs, we targeted splicing of exons that either disrupt the protein open-reading frame when skipped (exon 2) or that encode critical structural features of the protein such as the linker (exon 3), or LRR domains (exon 5-9) (Figure 1A). We designed and tested phosphorodiamidate morpholinos (PMO) ASOs that base-pair at the 3’ or 5’ splice site regions (ss) of exon 2, 3, 5, 6, 7, 8, and 9. PMOs were transfected into differentiated, macrophage-like THP-1 cells. ASO-induced skipping of exon 2 removes the canonical start codon and ASO-induced skipping of exons 3,5,6,7,8, and 9 is predicted to yield mRNAs with intact open-reading frames that encode protein isoforms with potentially altered functions. The effect of inducing the different NLRP3 isoforms and decreasing full-length NLRP3 using ASOs was measured by activating the inflammasome complex and measuring cytokine release. A non-targeted ASO (ASO-C) as a control for non-specific effects.

Cells were primed with lipopolysaccharide (LPS) and subsequently activated with ATP to initiate the NLRP3 inflammatory response. RNA and protein were collected from the cells and splicing was analyzed by reverse transcription PCR (RT-PCR) and sequencing of the resulting amplicons. Treatment with ASO-Δ2, targeting exon 2, resulted in splicing from an alternative 5’ss in the upstream untranslated region of exon 2 to the 3’ss of exon 3, which splices out the natural start codon of the transcript. ASO-Δ3, Δ5, Δ7, and Δ9 induced skipping of their respective exons. ASO-Δ6, targeting the 3’ss of the exon, activated an in-frame, cryptic 3’ss. ASO-Δ8 activated an out-of-frame 5’ss, resulting in partial exon 8 skipping and creation of a stop codon in exon 9 (Figure 1, A and B). Some ASOs also induced a very low level of additional splicing events. Overall, treatment of cells with ASOs targeting exons 2, 3, 5, 6, 7, and 8, caused a significant decrease in full-length *NLRP3* mRNA compared to ASO-C (Figure 1C).

NLRP3 protein from ASO-treated cells was analyzed by immunoblot analysis. The predicted change in molecular weight of the protein isoforms is relatively small (8-14 kDa reduction) relative to the size of the full-length protein (∼120 kDa), making it potentially difficult to visualize migration differences on the blot (Figure 1A). ASO-induced splicing of exon 2 resulted in the appearance of a lower band representing the protein isoform encoded by the induced mRNA (Figure 2A). Modulation of *NLRP3* exons 2, 5, 6, and 8 splicing also resulted in a significant decrease in NLRP3 protein abundance (Figure 2, A and B). The ASO-induced exon skipping of in-frame transcripts are expected to produce proteins with small in-frame deletions of the targeted exon coding sequence, which we predict would result in a small change in size but not a reduction in overall abundance of the protein unless the induced isoforms are unstable.

**Figure 2.**
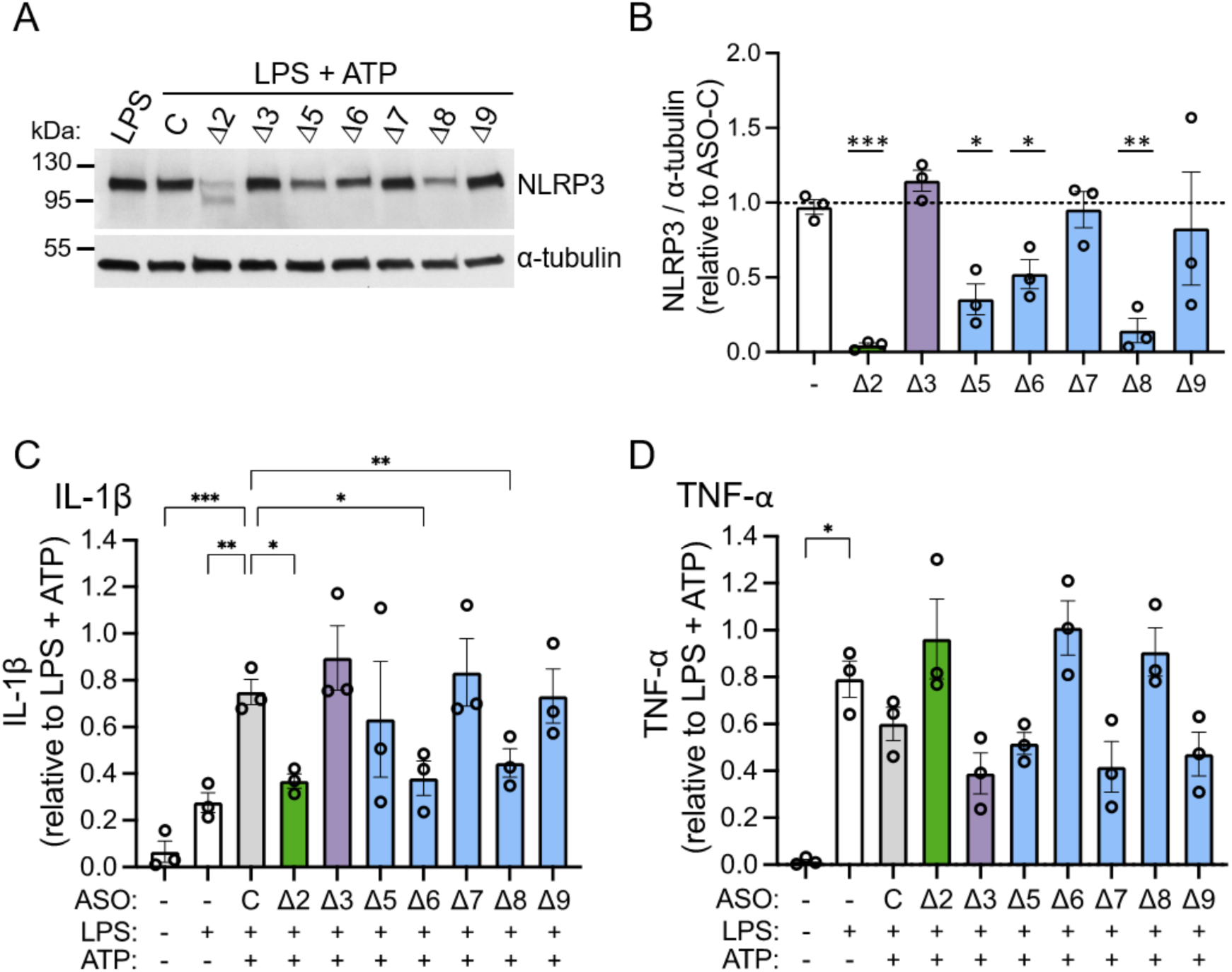
ASO-induced *NLRP3* alternative splicing inhibits inflammasome activation in human THP-1 cells. **(A)** Representative immunoblot of NLRP3 protein in THP-1 lysates treated with the indicated ASOs (40 µM), including a non-targeted ASO control (C), followed by activation with LPS and ATP or untreated and activated with LPS only. ⍺-tubulin was analyzed as a control. **(B)** Quantification of NLRP3 protein normalized to α-tubulin control. LPS only control (-) is included. Data presented as mean ± SEM; one sample t-test compared to non-targeted ASO control (ASO-C) set to 1, *P<0.05, **P<0.01, ***P<0.001; n=3. ELISA analysis of **(C)** IL-1β and **(D)** TNF-α released from THP-1 cells treated with ASOs prior to activation with LPS and ATP. Cytokine level was normalized to LPS/ATP stimulated cells. Data are presented as mean ± SEM and analyzed by repeated measures one-way ANOVA followed by Dunnett’s multiple comparisons test relative to non-targeted ASO control (C) for IL-1β and relative to LPS for TNF-α; *P<0.05, **P<0.01, ***P<0.001; n=3 independent experiments.

To evaluate whether ASO-mediated exon skipping and consequent modulation of NLRP3 protein expression reduced the cellular inflammatory response, IL-1β release into the media was quantified following activation. Treatment of cells with ASO-Δ2, ASO-Δ6, and ASO-Δ8 significantly reduced IL-1β secretion from THP-1 cells in response to LPS and ATP (Figure 2C). As expected, there were no changes in the secreted levels of tumor necrosis factor-α (TNF-α, TNF), a proinflammatory cytokine produced by activated macrophages independently of the NLRP3 pathway (Figure 2D).

The mechanism of NLRP3 inactivation varies for the different ASOs. ASO-Δ2 induces splicing out of the native translational initiation codon in exon 2 and thereby results in loss of NLRP3 translation. Protein modeling of NLRP3 Δ6 and Δ8 isoforms suggests that these deletions disrupt LRR-LRR interactions in the oligomerized form, likely explaining the reduced activity (Supplemental Figure 1).

Further testing of the most active ASOs, ASO-Δ2 and ASO-Δ6, identified their half-maximal inhibitory concentration (IC50) as 1.68 µM and 3.62 µM, respectively (Figure 3, A-E). Treatment of cells with these ASOs resulted in a dose-dependent decrease in IL-1β secretion following activation with LPS and ATP (Figure 3F). The level of inhibition of the inflammatory response was similar to that observed in cells treated with MCC950, a potent and specific small molecule inhibitor of NLRP3 (Figure 3G) (19). TNF-α levels were not decreased, suggesting that the ASOs are acting in an on-target manner specific to reducing NLRP3 activity without exerting an immunogenic effect (Figure 3F).

**Figure 3.**
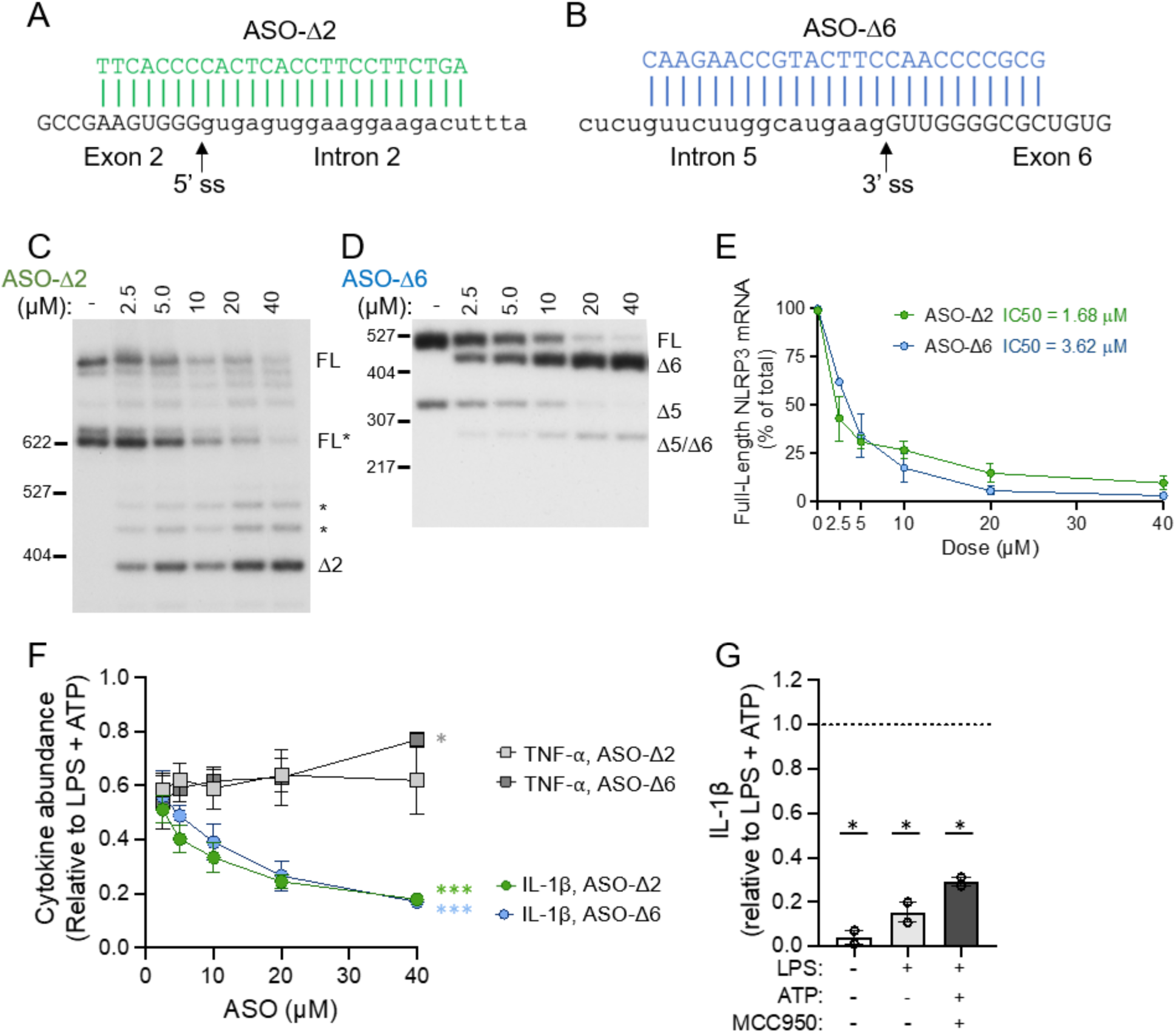
Dose-dependent activity of ASOs targeting *NLRP3* exon 2 and 6 splicing. Sequence alignment of (**A**) ASO-Δ2 and (**B**) ASO-Δ6 shown base-paired with their target sequence in *NLRP3.* The splice sites (ss) are indicated with the arrow. Exonic sequence is in capital letters and intronic sequence is in lowercase. **(C)** Representative radioactive RT-PCR analysis of *NLRP3* isoform expression in THP-1 cells transfected with increasing doses of (**C**) ASO-Δ2 or (**D**) ASO-Δ6, and 40 µM non-targeted ASO control (-). FL* denotes a naturally occurring isoform. Other potential spliced products indicated with asterisk (*). Products were amplified with primers specific to exon 1 and exon 4 in **C** and exons 4 and 7 in **D** and separated by PAGE. (**E**) RT-PCR quantification of full-length *NLRP3* mRNA relative to the predominant ASO-mediated isoforms. ASO potency was calculated using the half maximal inhibitory concentration (IC50) after plotting the data on a non-linear regression curve with a standard slope; ASO-Δ2 n=3; ASO-Δ6 n=1-2. **(F)** ELISA analysis of IL-1β and TNF-α released from THP-1 cells transfected with increasing doses (2.5 µM-40 µM) of ASO-Δ2 or ASO-Δ6 and activated with LPS and ATP. Cytokine levels were normalized to LPS and ATP stimulated cells. Slopes were calculated by linear regression analysis; *P<0.05, ***P<0.001; n=2-3 across 3 independent experiments. **(G)** ELISA analysis of IL-1β released from THP-1 cells treated with MCC950 and activated with LPS and ATP. Data are presented as mean ± SEM and analyzed by one sample t-test compared to LPS/ATP set to 1; *P<0.05; n=2 independent experiments.

To evaluate ASO-1′2 and NLRP31′2 in an in vitro model directly relevant to clinical, pathological NLRP3 activation, we treated human monocyte-derived macrophages (hMDMs) from individuals with CAPS harboring an NLRP3 p.L353P mutation with the ASO. Our results show that ASO-Δ2 modulates *NLRP3* exon 2 skipping in hMDMs, which correlates with a decrease in IL-1β release, confirming ASO activity in human primary cells and demonstrating its potential efficacy in blocking pathological NLRP3 activity (Supplemental Figure 2).

### ASO-mediated modulation of Nlrp3 splicing mitigates inflammasome signaling in immortalized mouse macrophages

To test the in vivo effects of inducing *NLRP3* spliced isoforms with ASOs, we needed to identify mouse-specific ASOs that induced skipping of the different *Nlrp3* exons because the human targeted *NLRP3* ASO target sequences are not highly conserved in the mouse sequence. For this, we designed mouse-specific ASOs that targeted the exon-intron junction of exon 2 (pyrin domain), 3 (linker domain), and 5-9 (LRR domains) and tested them in immortalized bone-marrow derived macrophages (iBMDMs) treated with LPS and ATP. ASOs were identified that induced skipping of exons 3, 5, 6, 8 and 9 and partial skipping of exon 2 (Figure 4, A and B). ASOs that induced skipping of these exons resulted in a significant decrease in full-length NLRP3 protein and/or the appearance of smaller proteins appeared on NLRP3 immunoblots, likely representing the protein isoforms encoded by mRNA with the targeted exon skipped (Figure 4, C and D). Treatment of cells with ASOs targeting the different exons also resulted in a significant decrease in cleaved CASP1 (p20) secretion (Figure 4, C and E). Likewise, IL-1β secretion was significantly reduced following treatment of cells with all ASOs except ASO-Δ7 (Figure 4F). Because inflammatory cell death occurs in iBMDMs in response to NLRP3 activation, we measured lactate dehydrogenase (LDH), a marker of pyroptosis, and found a significant decrease in levels following treatment of all ASOs except ASO-1′7, which did not effectively target exon 7 (Figure 4G). LPS-dependent TNF-α levels in the media were unaffected by *Nlrp3*-targeted ASO activity (Figure 4H).

**Figure 4.**
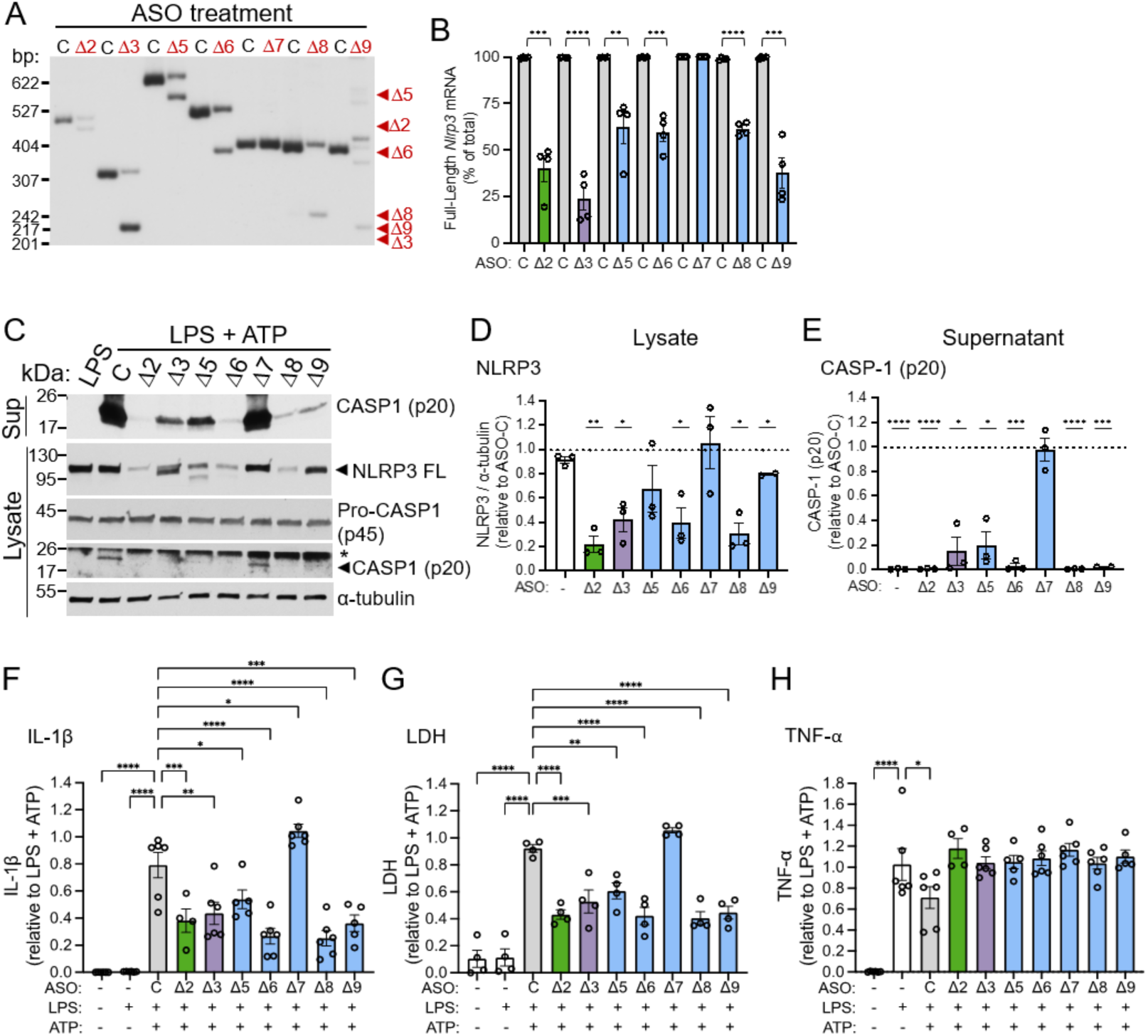
*Nlrp3*-targeted ASOs induce exon skipping and reduce inflammasome signaling in mouse iBMDMs. **(A)** Representative radioactive RT-PCR analysis of *Nlrp3* RNA expression in iBMDMs transfected with the indicated ASOs (40 µM), including a non-targeted ASO control (C) followed by activation with LPS and ATP. Products were amplified using specific primer sets for each exon and separated by PAGE. Red arrowheads indicate predominant ASO-induced spliced isoform for each ASO. Some ASOs also induced a low level of other alternatively spliced products. **(B)** Quantification of full-length *Nlrp3* mRNA relative to the predominant ASO-mediated isoforms. Data are presented as mean ± SEM, unpaired t-test compared to matched ASO control, **P<0.01, ***P<0.001, ****P<0.0001; n=4. **(C)** Representative immunoblot of NLRP3, Pro-CASP1 (p45) and cleaved CASP1 (p20) in iBMDM lysates and CASP1 (p20) in iBMDM supernatant treated with indicated ASOs (40 µM) or untreated, followed by LPS or LPS and ATP activation. ⍺-tubulin was analyzed as a control. Non-specific bands are denoted with asterisk (**\***). **(D)** Quantification of immunoblots of NLRP3 shown normalized to α-tubulin control. Data shown as mean ± SEM and analyzed by one sample t-test compared to non-targeted ASO control (ASO-C) set to 1, *P<0.05, **P<0.01; n=2-3. **(E)** Quantification of immunoblots of cleaved CASP1 (p20) secreted from iBMDM treated with ASOs and activated with LPS and ATP or untreated and activated with LPS, relative to ASO-C. Data are presented as mean ± SEM and analyzed by one sample t-test compared to ASO-C set to 1; *P<0.05, ***P<0.001, ****P<0.0001; n=2-3. **(F)** IL-1β, **(G)** LDH and **(H)** TNF-α released from iBMDMs untreated or treated with ASOs and activated with LPS or LPS and ATP. Cytokine levels were normalized to LPS and ATP stimulated cells. Data are presented as mean ± SEM and analyzed by one-way ANOVA followed by Dunnett’s multiple comparisons test relative to LPS for TNF-α and to ASO-C for IL-1β and LDH. n= 4-6; *P<0.05, **P<0.01, ***P<0.001, ****P<0.0001.

ASO-Δ2, one of the most active ASOs in iBMDMs, caused the use of an alternative 5’ splice site leading to an mRNA with a shifted open reading frame, creating a premature termination codon (PTC) in exon 3 (Figure 5, A and B). This ASO decreased full-length *Nlrp3* mRNA expression (IC50 of 16 µM) and IL-1β release in a dose-dependent manner (Figure 5, C and D).

**Figure 5.**
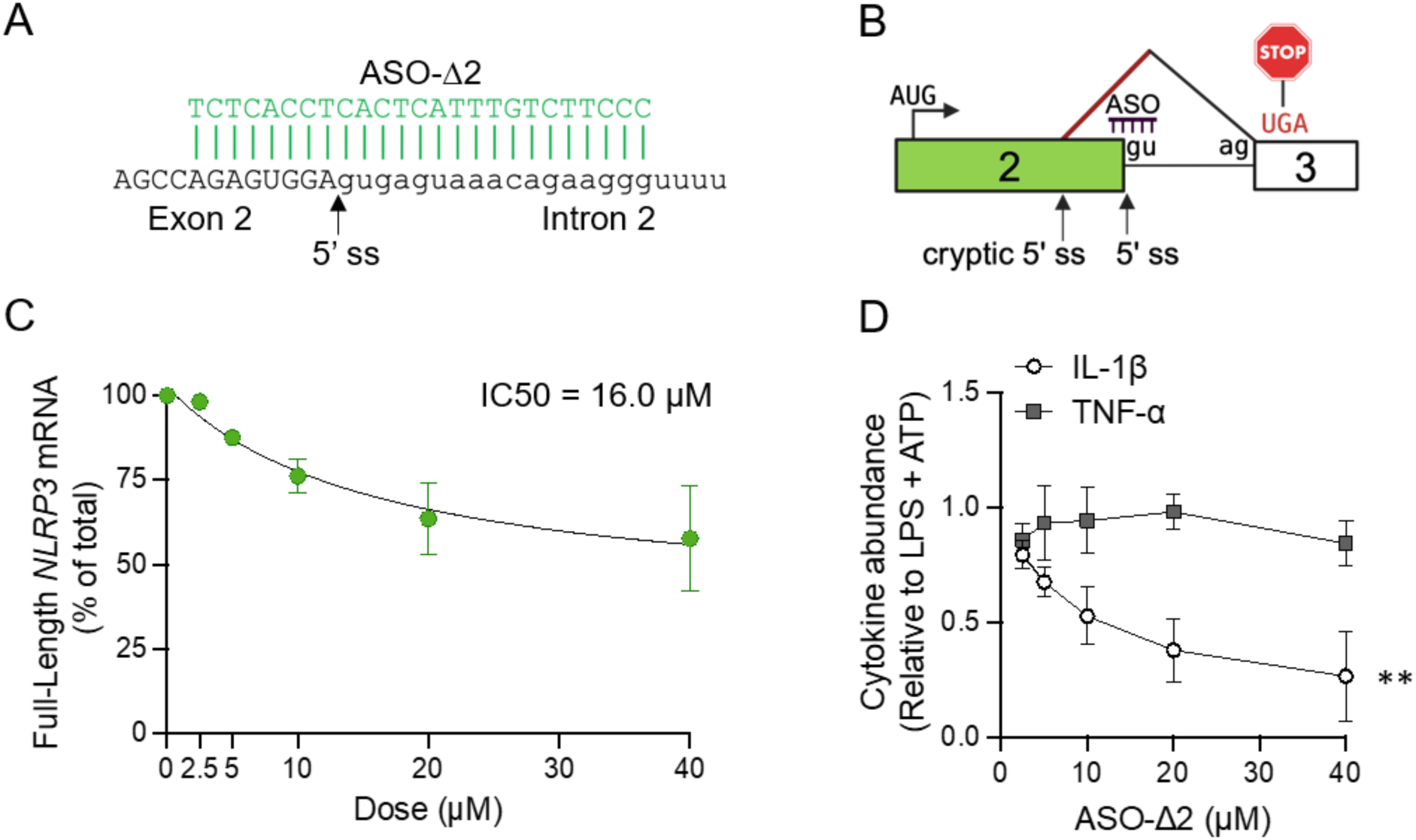
Dose-dependent modulation of *Nlrp3* exon 2 splicing by ASO-Δ2 in mouse iBMDM. **(A)** ASO-Δ2 shown base-paired to its target sequence at the *Nlrp3* exon 2 (capital letters)/intron 2 (lowercase) junction. The 5’ splice site (ss) of exon 2 is shown with an arrow. **(B)** Illustration of ASO-Δ2 mechanism of action showing activation of a cryptic 5’ ss in exon 2 that shifts the open reading frame and results in a premature termination codon in exon 3. Boxes are exons and lines are introns. ASO binding site is labeled. Diagonal black line shows splicing of introns. Red diagonal line indicates ASO-induced splicing event. AUG and UGA indicate the translational initiation and termination codon, respectively. **(C)** RT-PCR analysis of correctly spliced exons (full-length) from *Nlrp3* mRNA relative to the predominant ASO-mediated isoform from iBMDM cells transfected with increasing doses of ASO-Δ2 (2.5 µM-40 µM). ASO potency was determined using the half maximal inhibitory concentration (IC50) after fitting the data using a nonlinear regression curve with a standard slope; n=3 independent experiments. **(D)** ELISA analysis of IL-1β and TNF-α secretion from iBMDM cells transfected with increasing doses of ASO-Δ2 (2.5 µM-40 µM) and activated with LPS and ATP. Cytokine levels were normalized to LPS and ATP stimulated cells. Individual data points are presented as mean ± SEM. The slopes of the dose response curves were calculated by linear regression analysis, **P<0.01; n=3 independent experiments.

### *ASO-Δ2* suppresses systemic inflammation in an LPS model of sepsis

To examine the therapeutic potential of ASO-Δ2 in vivo, we tested its efficacy in a mouse model of acute inflammation that employs LPS, a potent pro-inflammatory endotoxin found in the outer membrane of gram-negative bacteria that activates innate immune signaling and thereby mimics sepsis, a life-threatening sequela of infection. Previous studies have shown that peripheral LPS injection induces an NLRP3-dependent inflammatory response in mice (20). To assess ASO-1′2 in this model, wildtype (WT) mice were treated with three doses of ASO-Δ2, ASO-C, or vehicle control by intraperitoneal (IP) injection on day 1, 3, and 6. On day 7, mice were injected with LPS or vehicle control and euthanized 3 hours later (Figure 6A). Plasma from the mice treated with ASO-Δ2 had significantly reduced IL-1β and interleukin-6 (IL-6), a downstream pro-inflammatory cytokine, but not TNF-α, compared to controls (Figure 6, B-D). These results demonstrate the specific anti-inflammatory activity of ASO-induced *Nlrp3* exon 2 skipping in vivo.

**Figure 6.**
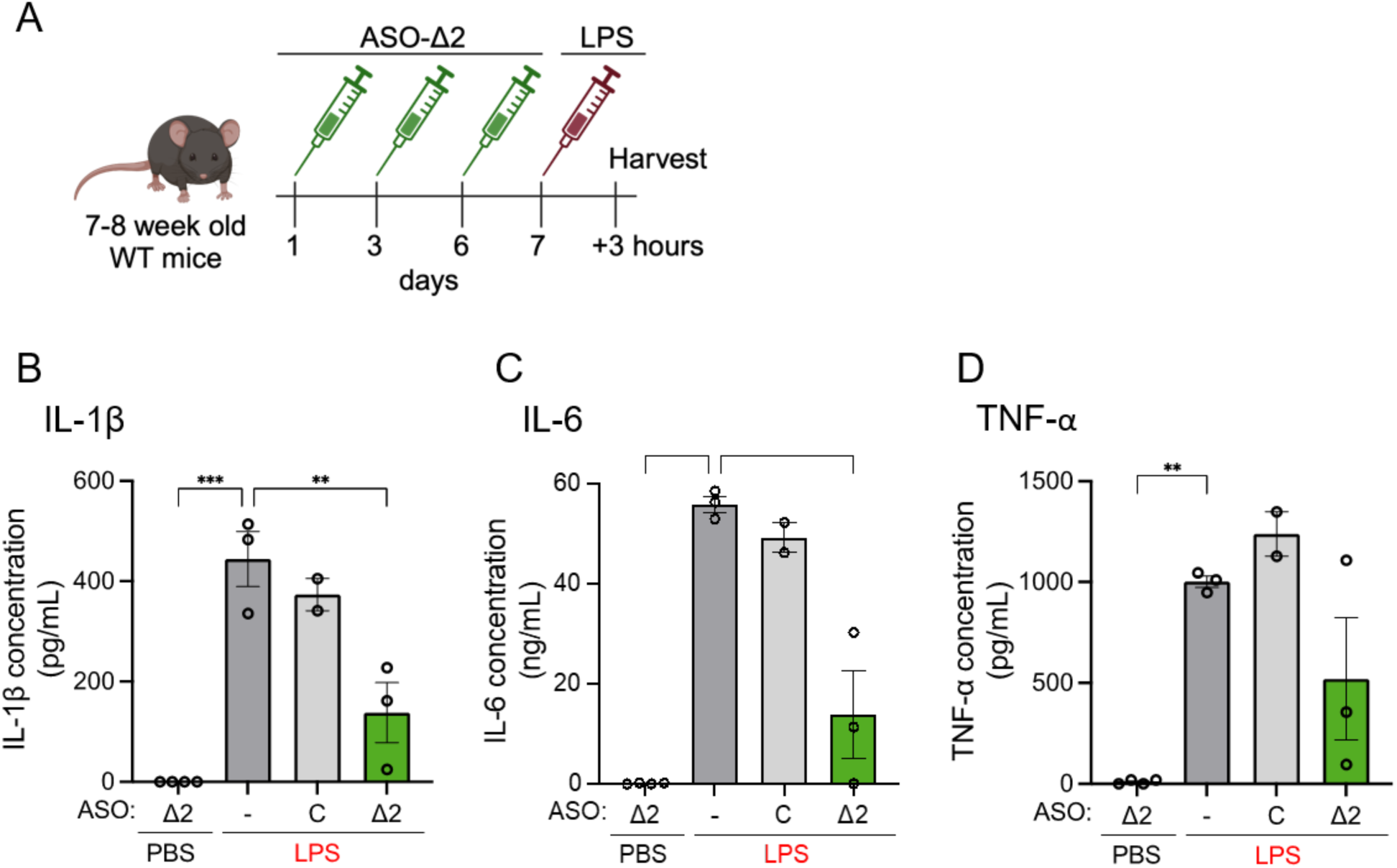
ASO-induced *Nlrp3* alternative exon 2 splicing reduces systemic inflammation in vivo. **(A)** Schematic of dosing regimen of 7-8 week-old female wildtype (WT) mice. Mice were treated three times with either PBS vehicle (n=3), non-targeted control ASO 100 mg/kg (n=2), or ASO-Δ2 100 mg/kg (n=3) followed by LPS or treated 3 times with ASO-Δ2 100 mg/kg (n=4) followed by PBS. ELISA analysis of **(B)** IL-1β, **(C)** IL-6, and **(D)** TNF-α in the plasma of the mice described in **A**. PBS vehicle (-) and non-targeted control ASO (C) were included as controls. Data are presented as mean ± SEM. Ordinary one-way ANOVA followed by Dunnett’s multiple comparisons test relative to PBS vehicle control; **P<0.01, ***P<0.001, ****P<0.0001.

### ASO treatment prolongs survival and alleviates inflammation in a CAPS mouse model

To further assess the therapeutic efficacy of ASO-Δ2 in a clinically relevant animal model, we investigated its activity in a mouse model of CAPS. *Nlrp3*^D301N/+ LysMCre+^ mice express constitutively active NLRP3 protein in the myeloid lineage. This mutation in mouse *Nlrp3* corresponds to the human D303N mutation which causes NOMID, a severe form of CAPS (21). *Nlrp3* ^D301N/+ LysMCre+^ mice develop severe multi-organ inflammation characterized by excessive secretion of proinflammatory cytokines, impaired growth, skin lesions, and perinatal death (21). The mice die prematurely likely due to dramatic cachexia and inability to meet the metabolic demands of systemic inflammation. *Nlrp3*^D301N/+ LysMCre+^ neonates were treated by subcutaneous injection with ASO-Δ2 or vehicle control on postnatal (P) day 1, 3, and every 3 days thereafter until P24 (Figure 7A). ASO-Δ2 treatment significantly prolonged survival (P<0.0001, Gehan-Breslow-Wilcoxon test), with 50% of mice surviving to P30 (maximum survival to P57) compared to controls (50% survival P15; maximum survival to P20) (Figure 7B). Unaffected, WT pups all survived to the completion of the study (Figure 7B). Body weight and spleen size were not significantly different between ASO-treated and control groups (Supplemental Figure 3). As rash is one of the earliest and most prominent manifestations of disease in people with CAPS, we visually and histologically assessed skin lesions in the mice, present mostly on the abdomen and neck, and observed lower neutrophil infiltration in skin of the ASO-1′2-treated mice compared to vehicle controls (Figure 7, C-E). In addition to skin inflammation, another hallmark of CAPS is end-organ systemic inflammation mediated by cytokine release in the blood, which is evident in the mice by the elevation of IL-1β and IL-18 but not IL-6 or TNF-α (Figure 7, F-I).

**Figure 7.**
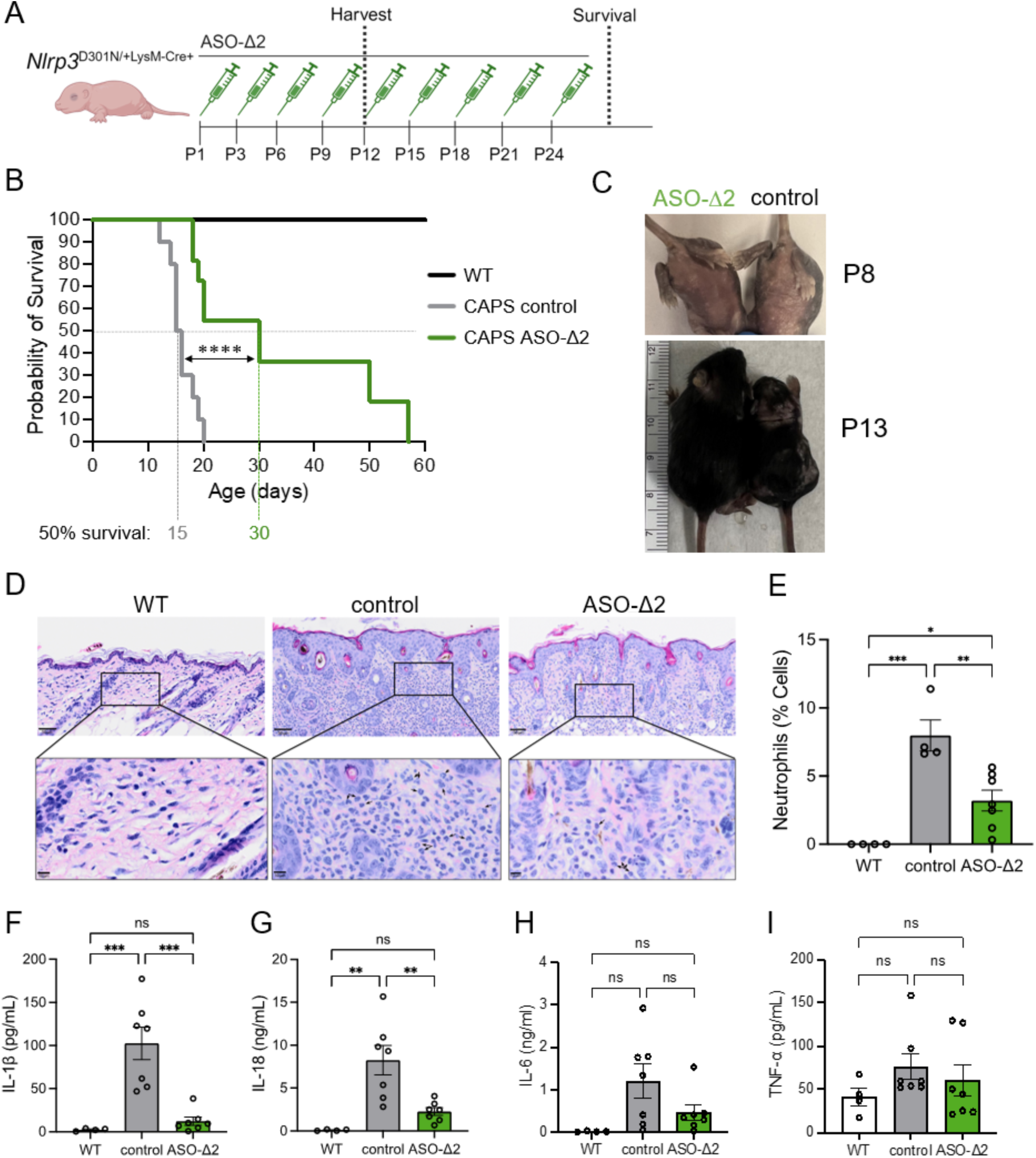
ASO-Δ2 prolongs survival and alleviates systemic inflammation in CAPS mice. **(A)** Schematic of dosing regimen in *Nlrp3^D301N/+ LysMCre+^* CAPS neonates treated with ASO-Δ2 or PBS control. Untreated wildtype, unaffected mice were used as controls. **(B)** Kaplan–Meier survival curve of mice treated with ASO-Δ2 (n=11), PBS control (n=11) and untreated wildtype (WT; n=12). The day that 50% of the mice for each group were alive is shown (50% survival). Curve comparison for ASO-Δ2 (n=11) vs. PBS control (n=11) Gehan-Breslow-Wilcoxon test; ****P<0.0001. **(C)** Images of representative ASO and control-treated *Nlrp3^D301N/+ LysMCre+^* mice at postnatal day 8 (P8) and P13. **(D)** Representative images of H&E staining of skin of ASO-1′2-treated *Nlrp3^D301N/+ LysMCre+^* (n=7), PBS control-treated (n=4), or untreated WT (n=4) mice. Scale bars, 50 μm top row and 10 μm in the magnified inset. Black arrows denote representative neutrophil morphology. **(E)** Quantification of neutrophil infiltration of skin from ASO-1′2-treated mice (n=7), PBS control-treated (n=4), and untreated WT (n=4) mice. One-way ANOVA with Tukey’s multiple comparisons test; *P<0.05, **P<0.01, ***P<0.001. ELISA analysis of **(F)** IL-1β, **(G)** IL-18, **(H**) IL-6, and **(I)** TNF-α levels in the serum of ASO-Δ2 (n=7), PBS control treated CAPS pups (n=7), and untreated WT pups (n=4) collected at P12. Data are mean ± SEM, one-way ANOVA followed by Tukey’s post-hoc analysis; **P<0.01, ***P<0.001, non-significant (ns).

ASO-Δ2-treated mice had levels of IL-1β and IL-18 similar to untreated, WT baseline expression, demonstrating that ASO-induced *Nlrp3* exon 2 skipping has a systemic anti-inflammatory, NLRP3-dependent effect in mice with a CAPS mutation associated with constitutive NLRP3 activation. ASO-induced *Nlrp3* exon 2 splicing in the liver confirms ASO activity in the treated mice (Supplemental Figure 3C). These data demonstrate that ASO-mediated modulation of *Nlrp3* exon 2 splicing ameliorates the clinical phenotype of CAPS in mice.

## Discussion

Inflammation is increasingly recognized as a contributing factor to the development of serious, often chronic diseases and cancers. Targeted reduction of pathological inflammation is an area of significant unmet therapeutic need. The NLRP3 inflammasome contributes to unwanted inflammation in numerous conditions. Direct NLRP3 inhibition is a means to short-circuit the inflammatory response early in the activation pathway to specifically control aberrant NLRP3-associated inflammation and lessen disease severity. Here, we demonstrate that splice-switching ASOs provide a specific and effective therapeutic strategy for suppressing NLRP3-mediated inflammatory signaling. These ASOs induce splicing alterations result in mRNAs encoding non-functional NLRP3 protein isoforms that are unstable and/or lack functional domains important for inflammasome assembly and activation.

Our systematic testing of ASO-induced skipping of seven of the ten NLRP3 coding exons revealed that an ASO that modulates exon 2 splicing is one of the most effective at down-regulating NLRP3 activity. The human *NLRP3* exon 2-targeted ASO blocks splicing at the predominant exon two 5’ splice site (donor site) and activates splicing at an alternative 5’ splice site in the exon, upstream of the start codon. Splicing at this upstream 5’ splice site to the native 3’splice site of exon 3, results in splicing out of the AUG start codon in exon 2, thereby decreasing NLRP3 protein expression due to the absence of a translation initiation site. ASO-Δ2 modulation of *NLRP3* exon 2 splicing decreases IL-1β release in hMDMs from patients with CAPS (Supplemental Figure 2), confirming ASO activity in human primary cells and offering promise for ASO-mediated exon skipping effectiveness in mitigating inflammasome signaling in disease-relevant cells. Further support for this conclusion comes from in vivo testing. An ASO targeted to mouse *Nlrp3* exon 2 resulted in the activation of a cryptic 5’ splice site in exon 2 upstream of the native site, which causes an open-reading frame shift and a premature termination codon (PTC), resulting in an mRNA that is predicted to make a small peptide (78 aa). The PTC also makes the transcript susceptible to nonsense-mediated mRNA decay (NMD) (Figure 5). Overall, we demonstrate that both human and mouse exon 2 targeted ASOs are effective at modulating splicing and reducing NLRP3.

The therapeutic potential of using ASO-1′2 to treat systemic inflammation and alleviate disease severity is supported by our results demonstrating efficacy in mouse models of pathological inflammation. Administration of ASO-Δ2 partially suppresses IL-1β and IL-6 levels in the plasma in response to peripheral LPS challenge, evidence of the protective effect against NLRP3 activation. Furthermore, ASO-1′2 reduced functional NLRP3 signaling in a mouse model of CAPS expressing a missense mutation in *Nlrp3* (D301N). ASO-1′2 prolonged survival and decreased systemic inflammation relative to controls, suggesting that ASO-mediated NLRP3 inhibition protects against constitutive inflammasome activation in vivo. Improved skin lesions, return of hair follicles, and reduction in neutrophil infiltration in the skin indicate potential efficacy of ASO-based approaches in addressing other inflammatory skin diseases. We did not detect ASO-induced *Nlrp3* exon skipping in whole blood collected on P12 possibly due to decreased stability of the PTC-containing *Nlrp3*1′ex2 mRNA in activated immune cells. A low level of *Nlrp3*1′ex2 was observed in the liver of mice treated with ASO-1′2, evidence of ASO-induced splicing in vivo (Supplemental Figure 3C). Together, these results show that ASO-mediated reduction in functional NLRP3 inflammasome signaling can therapeutically reduce disease-severity in mouse models of inflammation.

*NLRP3* exon 5 skipping has been identified by others as a naturally occurring splicing event as well, and the encoded protein has been shown to lack activity, which supports the hypothesis that alternative splicing may be a regulatory mechanism for responding to danger signals (22). We and others do not find an increase in exon 5 skipping in response to LPS, suggesting that these conditions do not induce alternative exon 5 splicing (23). Nonetheless, given that NLRP3 has naturally occurring isoforms, we propose that targeting splicing with ASOs is a means of harnessing the natural regulatory mechanism to generate an effective and safe therapeutic response.

Splice-switching ASOs targeting exons 6 or 8 splicing also result in impaired inflammasome signaling. Human ASO-1′6 activates an in-frame alternative 3’ splice site resulting in deletion of a small segment of the LRR domain (774-794 aa) and introduction of a single nucleotide polymorphism at position S795C. Human ASO-1′8 activates a cryptic 5’ splice site in exon 8 resulting in a shifted reading frame and creation of a PTC in exon 9, making the transcript susceptible to NMD. The functional impact that we observe experimentally by inducing an internal deletion and protein truncation with ASO-1′6 and 1′8, respectively, is supported by structural analysis of the protein isoforms as well. Recent cryogenic electron microscopy studies suggest that human NLRP3 forms an oligomer consisting of 10 subunits, creating a double cage-like structure (Supplemental Figure 1). Neighboring LRR domains from the two discs interact with each other front and back (24, 25). Our protein modeling of NLRP3 isoforms indicates that the Δ6 isoform lacks a region of the LRR involved in the convex interaction and that the Δ8 isoform deletes most of the regions making up the concave interactions. We predict that both isoforms, but particularly the Δ8 isoform, disrupt the stability of the decameric cage. The Δ6 isoform could both weaken the convex interactions as well as alter the concave interactions by changing the spacing of the LRR repeats C-terminal to the deletion. Importantly, a single ASO-induced NLRP3 isoform monomer can potentially destabilize an oligomerized structure formed with full-length NLRP3, thereby exerting a dominant negative effect that could amplify the effect of the ASO treatment. The LRR deletion in NLRP3 1′6 is unlikely to affect the formation of the decameric disk, as these interactions are mediated by the FIS and NACHT domains that are not predicted to be affected in the Δ6 and Δ8 isoform structure. In contrast, the Δ8 isoform lacks many of the amino acids involved in the full-length LRR-NEK7 interaction, which predicts a weakened interaction (Supplemental Figure 1). Mouse ASO-1′6 and 1′8 induce skipping of the entire targeted exon, which results in attenuation of inflammasome activation. The effect is likely due to deletions of segments of the LRR domain. Mouse NLRP3 subunits also form a dodecamer with interacting LRR domains to support complex stability (26). Our results suggest that ASO-mediated deletions of LRR segments may destabilize the 12-subunit structure, offering an explanation as to how these isoforms are less active.

An alternative ASO-based strategies to reduce NLRP3 has been previously reported. This approach involved an RNase H-targeting ASO (gapmer), which causes degradation of the RNA, a decrease in *Nlrp3* expression and extended the life of mice modeled to have CAPS (27). These results further support ASOs for modulating NLRP3 activity. Both splice-switching and RNase H-targeting ASOs have had clinical success in treating other diseases, making them promising candidate platforms for treating NLRP3-associated conditions (28–31).

The amount of ASO used in our in vitro studies is consistent with other PMOs used to target splicing of other transcripts in diseases such as Duchenne muscular dystrophy (32) and cystic fibrosis (33–35). Unlike other commonly used modifications such 2’MOE-PS, PMOs typically require higher doses in vitro likely due to their different chemical properties (positive vs neutral charge), delivery modalities (lipid reagent vs. Endoporter), the specific cell-type and other still unclear requirements for delivery to cells in culture. The doses used in vivo are consistent with those reported in the literature for approved therapies administered at doses as high as 30 mg/kg weekly (32, 36, 37).

ASOs offer an alternative to current treatment approaches and therapeutic strategies for targeting the NLRP3 inflammasome. To date, there are three IL-1 blockers approved by the FDA and EMA (11). However, because IL-1β is an important signaling component of other inflammasomes, not only the NLRP3 inflammasome, IL-1 inhibition compromises the innate immune system in a more global manner, thereby posing a high risk for infections and other potentially dangerous side-effects (38). NLRP3-specific inhibitors offer a more targeted approach to treating pathogenic inflammation mediated by the NLRP3 inflammasome and can also address other effects of NLRP3 activation, such as IL-18 secretion and pyroptosis, which contribute to lingering inflammation in IL-1β knockout CAPS mice (39). Recent studies have reported on small molecule inhibitors that target NLRP3 and alleviate inflammation associated with CAPS and other conditions (3, 19, 40, 41). These NLRP3 inhibitors are more effective than IL-1β inhibitors in mouse models of CAPS, suggesting that inactivating the inflammasome directly will provide greater therapeutic value than targeting downstream released cytokines such as IL-1β. Though promising results have been reported with NLRP3 inhibitors for a number of conditions, toxicities and off-target effects have been a concern in some cases and small molecule NLRP3 inhibitors are less effective for CAPS-associated mutations that disrupt the binding pocket of such compounds (42, 43). Hence, manipulating NLRP3 gene expression directly using ASOs is an attractive alternative approach for treating NLRP3-driven inflammation.

This study demonstrates a systematic approach for identifying potentially therapeutic splice-switching ASOs and specifically demonstrates the therapeutic potential of targeting *NLRP3* splicing with ASOs to control pathogenic inflammation. The identified ASOs offer a potentially therapeutic alternative to approved IL-1β drugs or small molecule NLRP3 inhibitors currently in clinical trials. The growing success of ASO drugs in the clinic and in clinical trials, supports the promise of an *NLRP3*-targeted ASO as a safe, durable, and highly specific therapeutic for treating NLRP3-mediated inflammatory disease.

## Author contributions

RK and MLH conceived and designed the study. RK, JO, JLC and EMV conducted the experiments. RK, JO, JLC, FJD and MLH analyzed the data. CDP performed protein modeling. RK and MLH wrote the original draft of the manuscript. RK, JO, JLC, EMV, CDP, HMH, FJD and MLH reviewed and edited the manuscript. HMH and MLH acquired funding and resources and supervised the project.

## Acknowledgments

The authors thank the patients for providing blood samples for this study, Drs. Venkat Magupalli and Hao Wu for the iBMDM cells and Dr. Reid Oldenburg for histopathological assessments. Some figure panels were created using BioRender.com.

This work has been funded by NIH R01 AG060195 and University discretionary funds to MLH and NIH R01 AI15586901 to HMH. UCSD School of Medicine Microscopy Core was supported by P30 NS047101.

Conflicts of interest: MLH has consulted with ProMIS Neurosciences and QurAlis and has had research collaborations with Ionis Pharmaceuticals; HMH has consulted for Novartis, Sobi, Ventyx, and Akros and has had research collaborations with Jecure, Zomagen, Takeda, and Inapill.

---

**Supplemental Figure 1. Modeling of NLRP31′ex6 and 1′ex8 proteins.**

The NLRP31′6 and 1′8 isoforms are predicted to affect the stability of the inactive decamer cage assembly and potentially the NLRP3-NEK7 interaction in the active disk assembly. **(A)** Comparison of the inactive decameric cage form of full-length NLRP3 (left, PDB id 7pzc; (21)) and Alphafold2 models of the Δ6 and Δ8 isoforms superimposed onto the full-length structure. **(B)** Depiction of the two unique interaction surfaces of the decameric NLRP3 cage assembly between the concave (blue) and convex (red) faces of the LRR domains (left). Mapping of the interactions onto a linear diagram of the NLRP3 proteins reveals that the Δ6 isoform specifically deletes a region of the LRR involved in the convex interaction (indicated green box) and that the Δ8 isoform deletes most of the regions making up the concave interactions (indicated green box). Both isoforms, but particularly the Δ8 isoform, would be predicted to disrupt the stability of the decameric cage. In the case of the Δ6 isoform, changes due to the deletion could both weaken the convex interactions as well as alter the concave interactions by changing the spacing of the LRR repeats C-terminal to the deletion. **(C)** Comparison of the active disk form of full-length NLRP3 (left, PDB id 8ej4; (42)) and Alphafold2 models of the Δ6 and Δ8 isoforms superimposed onto the full-length structure. **(D)** Depiction of the two unique interaction surfaces of the active decameric NLRP3 disk assembly between adjacent NLRP3 monomers (red) and NEK7 (blue) (left). Mapping of the interactions onto a linear diagram of the NLRP3 proteins suggests that both LRR deletions are unlikely to affect the decameric disk formation, as these interactions are mediated by the FIS and NACHT domains that are not affected in the Δ6 and Δ8 isoforms. In contrast, the Δ8 isoform eliminates many of the residues involved in the full-length LRR-NEK7 interaction (indicated green box), suggesting a weaken interaction.

**Supplemental Figure 2. ASO-Δ2 *NLRP3* exon 2 splicing in human monocyte-derived macrophages (hMDM) from persons with CAPS.**

**(A)** Radioactive RT-PCR analysis of *NLRP3* isoform expression in *NLRP3*^L353P/+^ hMDM cells transfected with 40 μM of ASO-Δ2 or non-targeted ASO control (C) for 24 hr prior to activation with LPS for 16 hr. Other potential spliced products are denoted with asterisk (*). FL* denotes a naturally occurring full-length isoform. Products were amplified using a forward primer specific for *NLRP3* exon 1 and a reverse primer in exon 4 and separated by PAGE. **(B)** Quantification of correctly spliced exons (full-length) from *NLRP3* mRNA relative to the predominant ASO-1′2 induced isoforms. Data are presented as mean ± SEM, unpaired t-test compared to matched control, *P<0.05; N=3 patients. **(C)** IL-1β secretion from hMDMs treated with ASO-Δ2 relative to non-targeted ASO (ASO-C). Data are presented as mean ratio ± SEM and analyzed by one sample t-test set to 1, *P<0.05; n=4 representing results of experiments using cells from three different patients, and cells from one patient analyzed in two independent experiments.

**Supplemental Figure 3. Analysis of weights and splicing in ASO-1′2-treated CAPS mice.**

**(A)** Growth curve of ASO-Δ2 and PBS control treated *Nlrp3*^D301N/+ LysMCre+^ mice and untreated wildtype (WT) mice. **(B)** Representative image of ASO-1′2-treated *Nlrp3*^D301N/+ LysMCre+^ and WT mice at postnatal day 21 (P21). **(C)** (top gel) Splicing analysis of full-length *Nlrp3* expression relative to partial Δ2 isoform in the liver using primers in exon 1 and 3. (middle gel) Amplification of the exon 2 skipped isoform only, using a primer that base pairs across the ASO-induced cryptic 5’ splice site in exon 2. (bottom gel) *β-actin* amplicon analyzed as a control of loading. Non-specific PCR products are denoted with asterisks (*). **(D)** Spleen size relative to total body weight of ASO-Δ2 and PBS control treated Nlrp3^D301N/+ LysMCre+^ pups at P12/P13 (n=7). Data are mean ± SEM, unpaired t-test.

